# Microfluidic Osteoarthritis-on-a-Chip for Evaluating Joint-Cell Responses to Tanezumab, a Humanized Anti-NGF Monoclonal Antibody

**DOI:** 10.64898/2026.07.13.738227

**Authors:** Hosein Mirazi, Scott T. Wood

**Affiliations:** Department of Nanoscience and Biomedical Engineering, South Dakota School of Mines and Technology, Rapid City, South Dakota, USA; Portland Laboratory for Biotechnology and Health Sciences, University of New England, Portland, Maine, USA; Department of Biomedical Sciences, University of New England, Biddeford, Maine, USA

**Keywords:** Osteoarthritis, Joint-on-a-chip, Microfluidic co-culture, NGF inhibition (tanezumab), Drug safety assessment, Predictive *in vitro* modeling

## Abstract

Osteoarthritis (OA) drug development remains constrained by preclinical models that fail to recapitulate the multicellular interactions that regulate human joint inflammation and extracellular matrix degeneration in response to investigational drugs. Tanezumab, a humanized anti-nerve growth factor monoclonal antibody developed for non-opioid pain relief, advanced to late-stage clinical trials but was discontinued due to unresolved joint-localized safety concerns, including rapidly progressive OA. This study evaluated whether a human microfluidic joint-on-a-chip co-culture system could detect early biomarker responses to tanezumab exposure that were not apparent in conventional chondrocyte monoculture.

Tanezumab was first tested in human chondrocyte monoculture under untreated and disease-like (i.e., IL-1β-treated) conditions. Across a 20-analyte panel of inflammatory and matrix-remodeling biomarkers, statistically significant monoculture responses to tanezumab were limited to decreased IL-1β from 335 to 132 pg/mL (∼0.39-fold) and increased IL-8 from 575 to 675 pg/mL (∼1.17-fold). Major OA-associated matrix-remodeling markers, including MMP-1, MMP-3, and MMP-13, remained largely unchanged, indicating that monoculture conditions are insufficiently sensitive to detect clinically predictive drug-related molecular changes.

Tanezumab was then evaluated in co-cultures containing chondrocytes, osteoblasts, fibroblast-like cells, and macrophages under low-inflammation (i.e., M0 macrophage-based) and high-inflammation (i.e., M1 macrophage-based) conditions. In the M0-based co-culture, tanezumab increased MMP-1 from ∼4.20 × 10^4^ to ∼6.20 × 10^4^ pg/mL (∼1.48-fold), MMP-3 from ∼8.00 × 10^4^ to ∼1.20 × 10^5^ pg/mL (∼1.50-fold), and MCP-1 from 2.85 × 10^3^ to 4.31 × 10^3^ pg/mL (∼1.51-fold). In contrast, the M1-based co-culture showed decreases in MMP-13 from ∼1.66 × 10^4^ to ∼1.17 × 10^4^ pg/mL (∼0.70-fold) and IFN-γ from ∼1.95 × 10^4^ to ∼1.56 × 10^4^ pg/mL (∼0.80-fold), changes that may appear beneficial despite the drug’s known clinical risks.

Collectively, these findings show that low-inflammation multicellular co-culture revealed coordinated matrix remodeling and inflammatory responses to NGF blockade that were missed in monoculture and were partly obscured in highly stimulated disease-like conditions. This platform may provide a useful, human-relevant approach for safety signal assessment and early evaluation of OA therapeutics within a defined context of use focused on joint-specific, tissue-level drug-response testing.

## 2 Introduction

Osteoarthritis (OA) is a prevalent degenerative joint disease and a leading cause of chronic pain and disability worldwide. ("Global, regional, and national burden of osteoarthritis, 1990-2020 and projections to 2050: a systematic analysis for the Global Burden of Disease Study 2021," 2023; Wu et al., 2025) Despite extensive research, no disease-modifying OA drug (DMOAD) has yet received regulatory approval. (Li et al., 2023; Oo & Hunter, 2022; Qvist et al., 2008) Thus, current pharmacologic treatments primarily manage symptoms, i.e., pain. This has driven the pursuit of safer, more effective, non-opioid pain therapies that can address chronic OA pain without long-term risks.

Among these strategies, inhibition of nerve growth factor (NGF), a member of the neurotrophin family of growth factors, was proposed to reduce pain signaling at the peripheral level. (Lane et al., 2010; McMahon et al., 1995; Schmelz et al., 2019; Woolf et al., 1994; Zhao et al., 2024) Tanezumab, a humanized monoclonal antibody targeting NGF (Fig. 1), was developed by Pfizer and Eli Lilly as a novel non-opioid analgesic. (Danehy, 2017; Wadhwa et al., 2019) Although clinical trials demonstrated improvements in pain and physical function, regulatory agencies raised safety concerns, particularly reports of rapidly progressive osteoarthritis (RPOA), which called into question the long-term risk-benefit profile. (Schmelz et al., 2019) Despite extensive clinical evaluation, including positive findings from Phase III trials, tanezumab development was ultimately discontinued after the FDA raised unresolved safety concerns. (May, 2021; McKenzie, 2021)

**Figure 1.**
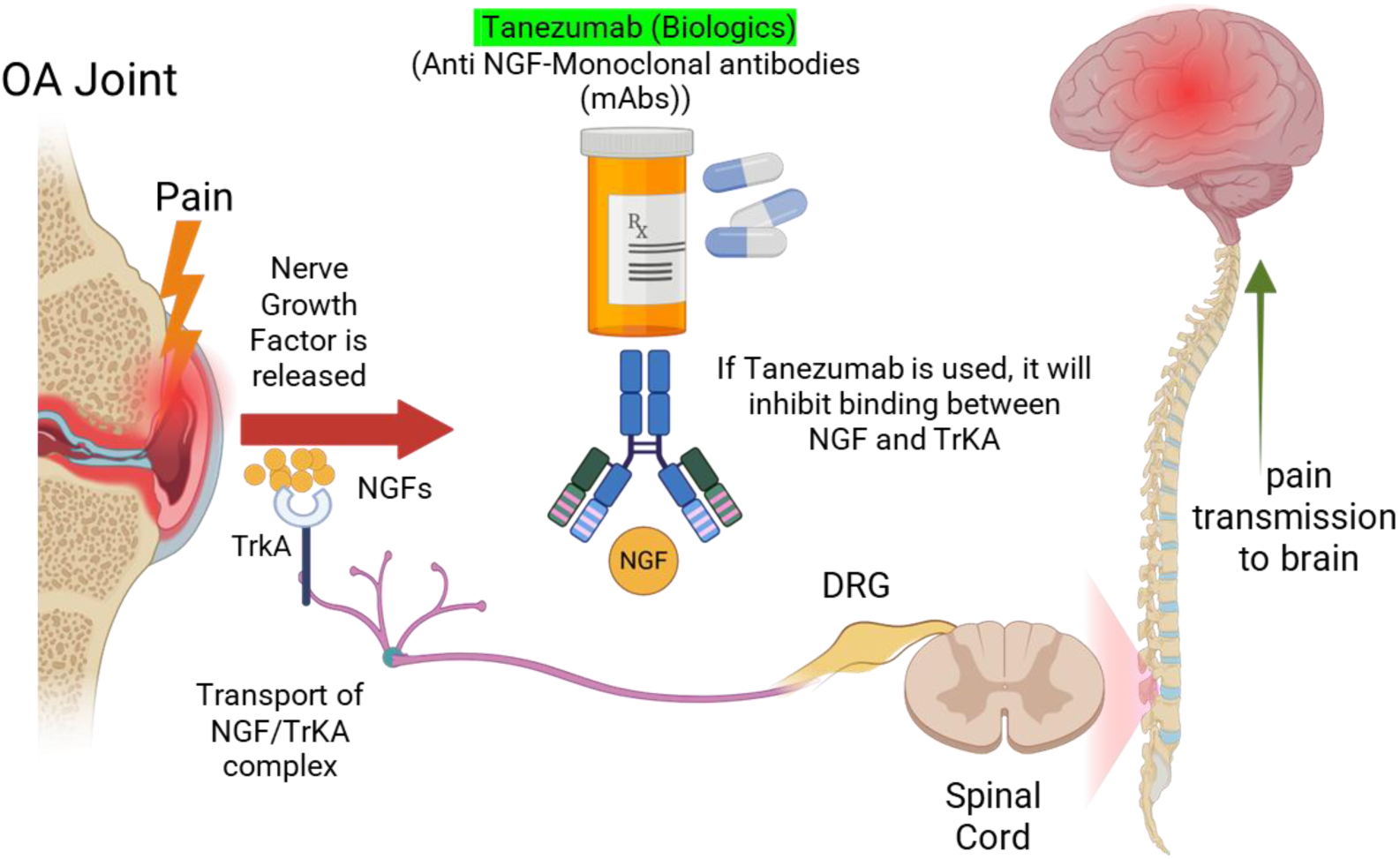
Tanezumab inhibition of NGF-mediated pain signaling in osteoarthritis. Schematic showing NGF release from the inflamed osteoarthritic joint, NGF binding to the TrkA receptor on peripheral sensory neurons, retrograde transport of the NGF/TrkA complex to the dorsal root ganglion, and subsequent pain transmission to the spinal cord and brain. Tanezumab blocks NGF-TrkA interaction, preventing peripheral sensitization and reducing nociceptive signaling. Created in BioRender. Mirazi, H. (2025) https://BioRender.com/fbwfcu7

The mechanisms underlying tanezumab-associated RPOA have not been fully established. Joint abnormalities observed early in the clinical program were initially suspected to represent osteonecrosis, prompting the FDA to place a partial clinical hold on tanezumab studies in 2010. (Hochberg, 2015; Pfizer, 2010) However, subsequent blinded adjudication determined that tanezumab was not associated with an increased incidence of primary osteonecrosis and instead identified RPOA as the predominant joint-safety phenotype. (Hochberg et al., 2016) Analgesic or neuropathic arthropathy has subsequently been proposed as one possible explanation. Under this model, pain relief reduces protective limitation of joint use, allowing greater loading of an already structurally compromised joint. (Roemer et al., 2023; Tiseo et al., 2014; Zhang et al., 2021) Supporting this hypothesis, tanezumab treatment in a rat medial meniscal tear (MMT) model prevented acute injury-associated gait deficiency and increased voluntary loading of the operated limb, resulting in greater cartilage damage. Delaying treatment until after the acute gait abnormality had resolved eliminated this effect. (LaBranche et al., 2017) The increased incidence of joint-safety events observed with tanezumab-NSAID combination treatment is also consistent with a contribution from analgesia-associated loading, although it does not distinguish increased loading from other pharmacologic or tissue-level interactions. (Hochberg, 2015; Hochberg et al., 2016)

Nevertheless, analgesia-driven overuse may not fully explain the clinical RPOA phenotype. The rat MMT model reproduced loading-associated cartilage damage but did not reproduce the hypotrophic bone destruction observed in patients with tanezumab-associated RPOA. (LaBranche et al., 2017) Multiple reviews have therefore concluded that the pathophysiology of tanezumab-associated RPOA remains unresolved and that additional joint- or tissue-level mechanisms may contribute. (Dietz et al., 2021; Schmelz et al., 2019; Zhao et al., 2022)

Importantly, NGF signaling is not restricted to nociceptive neurons. Human articular chondrocytes express NGF and its high-affinity receptor, TrkA. (Iannone et al., 2002) More recent preclinical evidence indicates that signaling through the neurotrophin receptor NGFR/p75NTR can regulate osteochondral inflammation and remodeling by limiting NF-κB activation, maintaining BMP-SMAD signaling and suppressing RANKL-associated bone resorption. (Zhao et al., 2024) Although these findings do not directly establish a mechanism for tanezumab-associated RPOA, they demonstrate that NGF signaling can influence joint-tissue biology independently of nociception. Additionally, synovial fibroblasts also express both NGF and TrkA, both of which have been shown to be upregulated in response to stimulation by the pro-inflammatory cytokines tumor necrosis factor-alpha (TNF-α) and interleukin-1beta (IL-1β). (Raychaudhuri & Raychaudhuri, 2009) These observations suggest that NGF blockade may alter coordinated responses among cartilage, bone, and synovial cells in ways not captured by pain measurements alone.

Thus, the case of tanezumab highlights a broader limitation in OA drug development: that conventional preclinical systems may not predict adverse outcomes arising from interactions among multiple joint tissues. Species differences in target biology, antibody pharmacology, joint structure, and mechanical loading can limit the translation of animal studies, while monoculture systems do not reproduce the intercellular signaling that coordinates inflammation, matrix remodeling, and bone-cartilage homeostasis.

In prior work and in a companion manuscript submitted concurrently to *Biofabrication*, we established and characterized a microfluidic co-culture model integrating chondrocytes, osteoblasts, fibroblasts, and macrophages within a shared recirculating microenvironment that supports paracrine communication. (Mirazi & Wood, 2025; Mirazi & Wood, 2026) The companion manuscript specifically evaluates the healthy M0-based and disease-like M1-based inflammatory configurations of this platform, whereas the present study uses these configurations to determine whether multicellular joint models improve the detection of tanezumab-induced biomarker responses. Each co-culture configuration was evaluated relative to an appropriately matched chondrocyte monoculture condition: an untreated monoculture for the healthy model and an IL-1β-treated monoculture for the disease model. Because the co-culture system enables communication among immune, stromal, cartilage, and bone-associated cells rather than modeling disease solely through direct cytokine stimulation of chondrocytes, it may provide a more physiologically relevant setting for evaluating drug-induced responses. (Mirazi & Wood, 2026)

This study addressed two related aims. The first was to determine whether human chondrocyte monoculture was sufficient to detect early biomarker responses potentially relevant to tanezumab-associated joint toxicity, including changes in the secretion of the OA-associated catabolic enzymes matrix metalloproteinases (MMPs) MMP-1, MMP-3, and MMP-13. We hypothesized that monoculture would exhibit a limited response because it does not capture the multicellular interactions present within the joint microenvironment.

The second aim was to determine whether the healthy or disease co-culture configuration detected biomarker responses to tanezumab that were absent from, or less apparent in, chondrocyte monoculture. We hypothesized that the co-culture models would reveal more RPOA-like changes in matrix remodeling and inflammatory biomarkers because they recapitulate paracrine communication among multiple joint-associated cell populations.

Together, these aims provide a context-of-use framework for evaluating whether a multicellular, human microphysiological system can improve the detection of drug-associated joint safety signals. By comparing tanezumab responses across monoculture and healthy and disease co-culture conditions, this study evaluates the potential utility of multicellular joint models for identifying adverse tissue-level responses that may be missed by conventional monoculture assays.

## 3 Materials and Methods

All microfluidic chip preparation, cell-handling workflows, seeding procedures, and media formulations followed the approaches established in our previous joint-on-a-chip studies. (Mirazi & Wood, 2025; Mirazi & Wood, 2026) Briefly, primary human osteoblasts (HOBs), primary human chondrocytes (HCHs), primary adult human dermal fibroblasts (HDFs), and THP-1 monocytes (derived from the peripheral blood of a 1-year-old male with acute monocytic leukemia) were obtained from commercial suppliers (PromoCell and ATCC) and cultured following previously established protocols, as shown in Fig. 2, and maintained in their respective growth media at 37 °C in a humidified atmosphere with 5% CO2. All adherent cell types were used at passages 2-5, with THP-1 monocytes cultured under standard suspension conditions.

**Figure 2.**
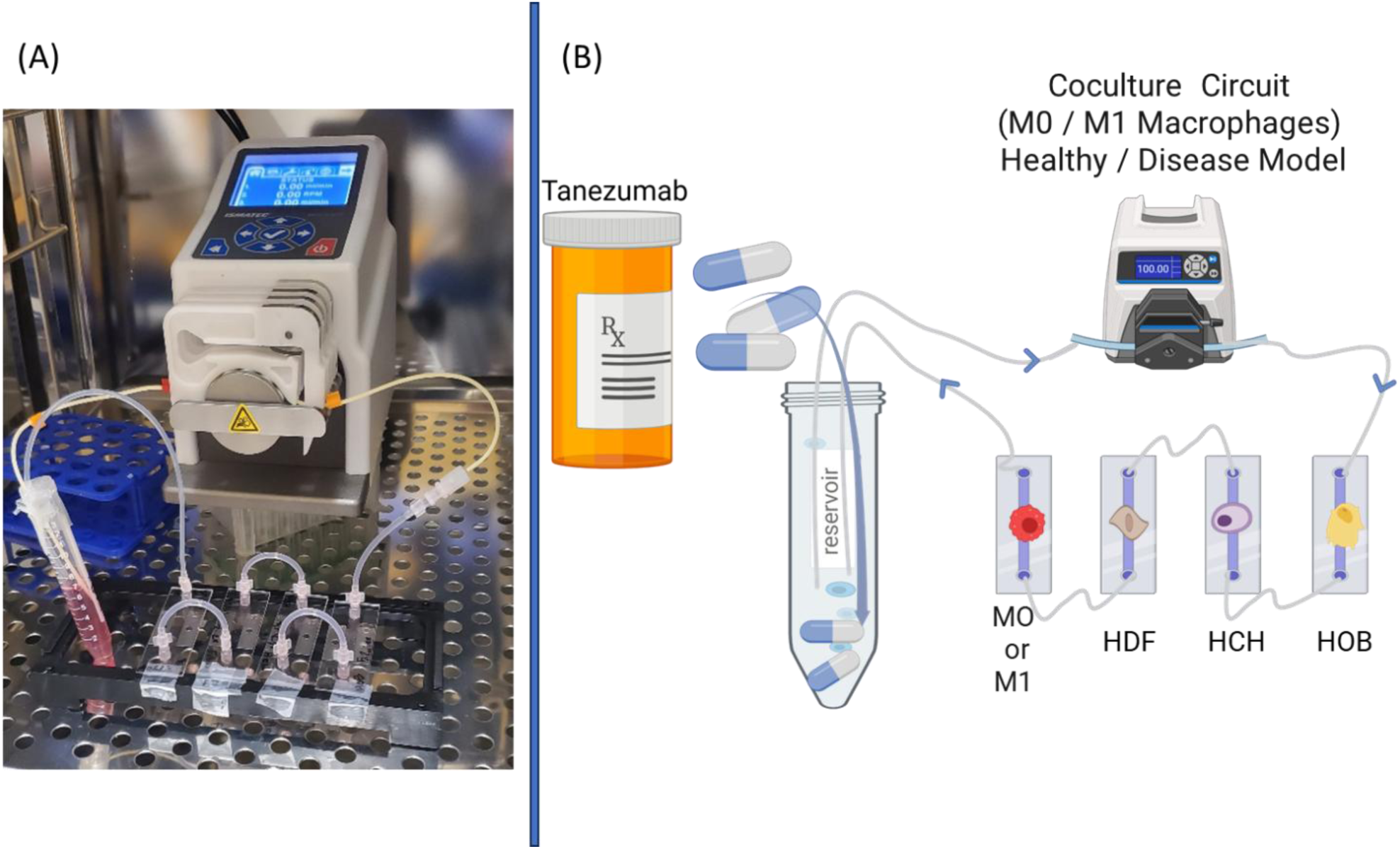
Experimental setup and schematic representation of tanezumab delivery in the microfluidic joint co-culture system. **(A)** Photograph of the microfluidic co-culture experiment showing Ibidi µ-Slide chambers seeded with human dermal fibroblasts (HDF), human chondrocytes (HCH), human osteoblasts (HOB), and macrophages connected in series to a peristaltic pump for continuous perfusion. Macrophages were included as either M0 (healthy-like) or M1 (disease-like) phenotypes**. (B)** Schematic illustration of the corresponding co-culture circuit, in which tanezumab is introduced into the shared reservoir and circulated through the interconnected microfluidic chambers to evaluate drug effects under healthy (M0 macrophages) and disease-like (M1 macrophages) inflammatory conditions. Created in BioRender. Mirazi, H. (2025) https://BioRender.com/guyc8ly

For the monoculture comparison, primary human articular chondrocytes were seeded under four experimental conditions: untreated baseline, tanezumab treatment, IL-1β stimulation to establish a disease-like inflammatory condition, and IL-1β-stimulated cultures that were subsequently treated with tanezumab. For the co-culture comparison (Fig. 2), microfluidic joint-on-a-chip models were generated under four conditions: healthy co-culture, healthy co-culture treated with tanezumab, disease-like OA co-culture, and disease-like OA co-culture followed by tanezumab treatment.

The untreated healthy and IL-1β-stimulated monoculture conditions, as well as the untreated M0-and M1-based co-culture conditions, overlap with reference/control conditions reported in a companion manuscript submitted concurrently to *Biofabrication*. (Mirazi & Wood, 2026) In that companion manuscript, these conditions are analyzed to characterize macrophage-mediated joint inflammation and paracrine crosstalk in the microfluidic co-culture platform. In the present study, the same reference conditions serve as baseline comparators for evaluating tanezumab-induced biomarker responses. Thus, although selected control data are shared between the two manuscripts, the studies address distinct hypotheses, use different treatment comparisons, and support different conclusions.

Tanezumab was purchased from MedChemExpress (MCE), prepared according to the manufacturer’s instructions, and dosing was adapted based on the exposure ranges reported in clinical trial NCT00809783. (*Extension Study Of Tanezumab In Osteoarthritis*, 2008) Tanezumab at 10 μg/mL, or equivalent volume PBS as the vehicle control, was added either directly to the chondrocyte monocultures or to the main reservoir of the microfluidic co-culture system to ensure uniform distribution across all cell compartments. Following drug administration, microfluidic cultures were subjected to 3 hours of controlled recirculation, then maintained under static conditions for approximately 36 hours to allow paracrine communication. Conditioned media were then collected from the chondrocyte compartment for biomarker quantification. All experiments were conducted in biological quadruplicate (n = 4), with identical media composition and incubation parameters applied across monoculture and co-culture conditions to ensure direct comparability.

To characterize drug-induced changes in inflammatory signaling and matrix-remodeling activity, conditioned media collected from the HCH compartments were analyzed using Quantibody® multiplex sandwich ELISA arrays (RayBiotech, Peachtree Corners, GA, USA). Two arrays were used in this study. The Human MMP Array Q1 (Cat. #QAH-MMP) was used to quantify MMP-1, MMP-2, MMP-3, MMP-8, MMP-9, MMP-10, and MMP-13, together with TIMP-1, TIMP-2, and TIMP-4. The Human Inflammation Array Q1 (Cat. #QAH-INF) was used to measure inflammatory cytokines and regulatory mediators, including IL-1α, IL-1β, IL-4, IL-6, IL-8, IL-10, IL-13, TNF-α, IFN-γ, and MCP-1. These biomarkers were selected to assess osteoarthritis-relevant changes in extracellular matrix turnover and inflammatory activity following tanezumab exposure.

All assays were performed according to the manufacturer’s instructions. After sample processing, the slides were submitted to RayBiotech for fluorescence scanning and data analysis. Fluorescence was detected using an InnoScan 710AL microarray scanner (Cy3 channel; Inopsys, Carbonne, France), and signal intensities were extracted and normalized using RayBiotech analysis software. Each analyte was printed in quadruplicate technical replicates on the array, and the median value of the four spots was used for each biological replicate. The resulting concentrations were used for comparisons across treatment groups to evaluate biomarker responses associated with tanezumab exposure.

All statistical analyses were conducted using GraphPad Prism version 10.3.1 (GraphPad Software, San Diego, CA, USA). Comparisons between two groups were performed using an unpaired two-tailed Welch’s t-test when appropriate. For experiments involving two independent variables, data were analyzed using ordinary two-way analysis of variance (ANOVA), followed by Fisher’s least significant difference (LSD) post hoc test using a single pooled variance. A *p*-value less than 0.05 was considered statistically significant.

## 4 Results

As shown in Fig. 3, multiplex fluorescence ELISA arrays were used to quantify cytokine and matrix-remodeling markers in conditioned media collected from the HCH regions of each model. Representative fluorescent scans from the Human Inflammation and Human MMP panels are shown in Fig. 3A-B, with each analyte printed in quadruplicate. Distinct differences in spot intensity were observed across monoculture and co-culture samples, reflecting condition-dependent variation in inflammatory signaling and extracellular matrix regulatory activity. These array outputs formed the basis for subsequent quantitative analyses of individual cytokines, MMPs, and related inhibitors across all experimental groups.

**Figure 3.**
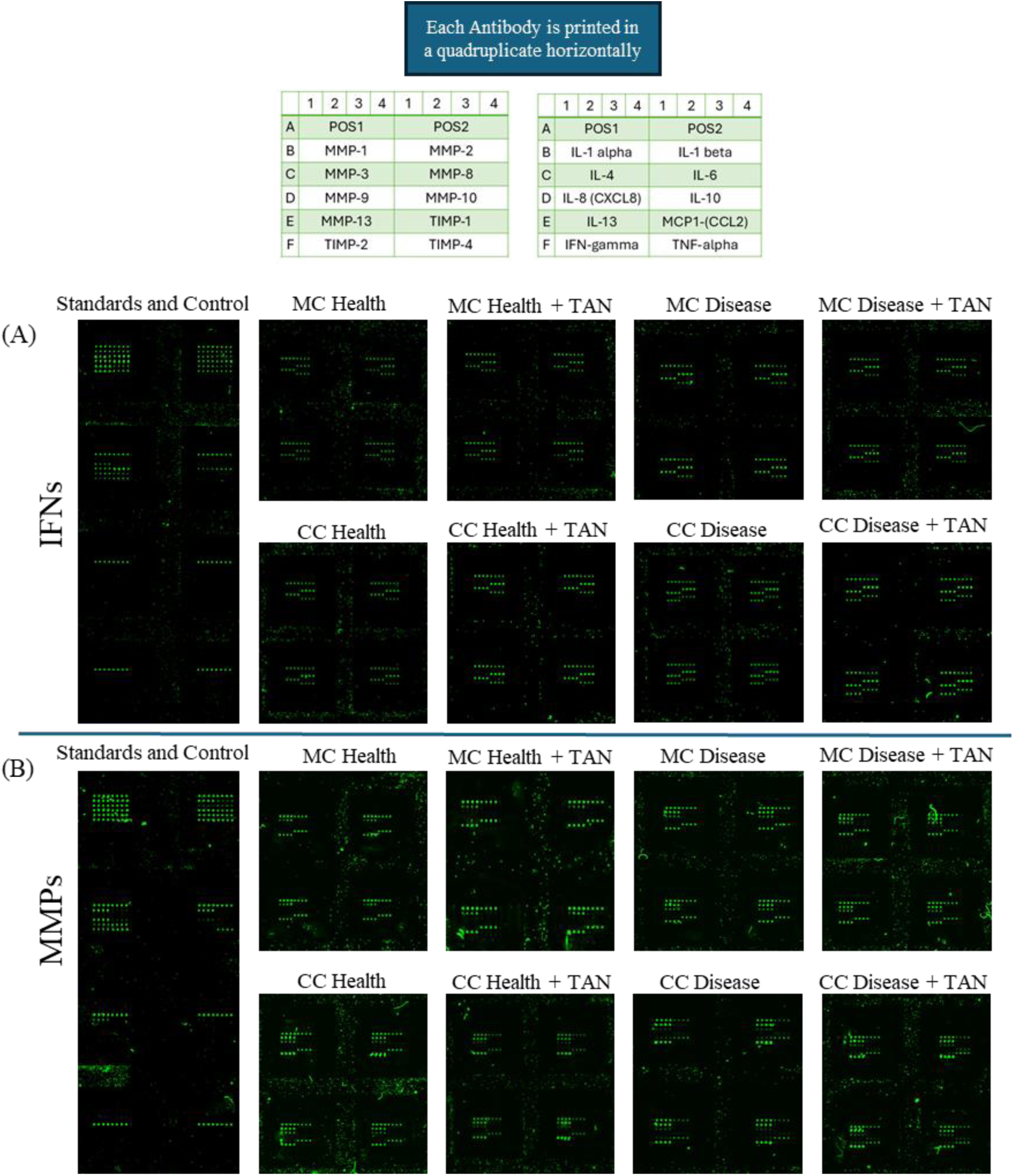
Representative multiplex fluorescence array scans. (A) Human Inflammation array and (B) Human MMP array, each printed in quadruplicate for quantitative reliability. Conditioned media from monoculture (MC) and co-culture (CC) systems were applied to assess tanezumab-induced changes in cytokines and matrix-remodeling markers. Differences in spot intensity reflect condition-dependent variations that were used for downstream quantitative analysis.

### 4.1 Monoculture Response to Tanezumab

As shown in Fig. 4, comparison between vehicle (untreated) and tanezumab-treated monocultures revealed significant differences in only two analytes, IL-1β and IL-8. All other measured markers remained largely comparable between groups.

**Figure 4.**
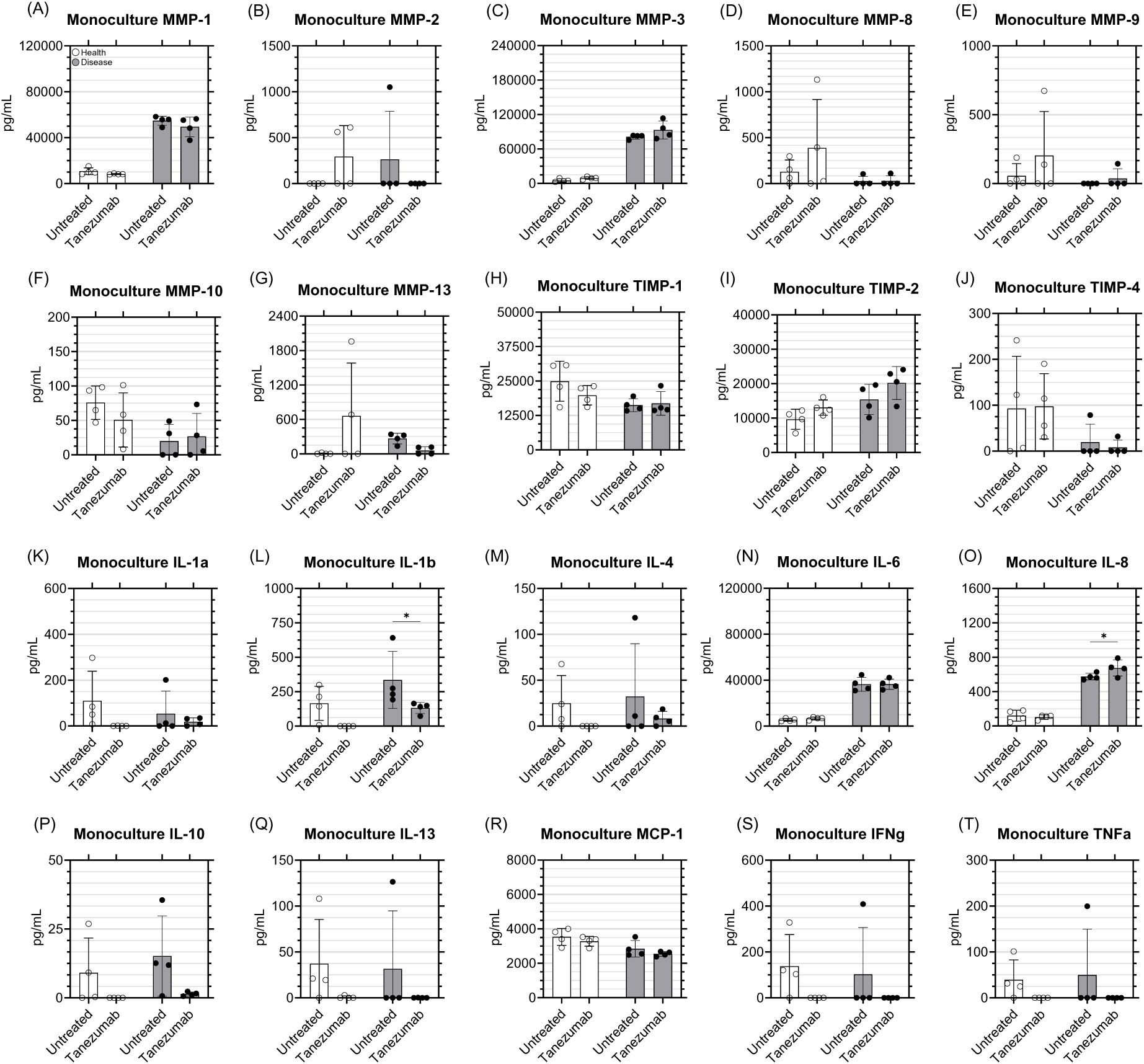
Quantification of inflammatory and matrix-remodeling markers in human chondrocyte monoculture following tanezumab treatment. Multiplex fluorescence ELISA measurements of cytokines, chemokines, and MMP/TIMP family members were obtained from conditioned media of four monoculture conditions: untreated healthy, healthy + tanezumab, IL-1β-stimulated disease-like, and IL-1β + tanezumab. Bars represent mean ± SD (n = 4). Among all analytes, only IL-1β and IL-8 exhibited statistically significant differences between untreated and tanezumab-treated disease-like cultures (p < 0.05), whereas no significant changes were detected in the healthy monoculture.

For IL-1β (Fig. 4L), disease-like monoculture showed lower concentrations following tanezumab treatment, decreasing from 335 pg/mL in untreated cultures to 132 pg/mL after exposure (p < 0.05). In contrast, healthy monoculture showed no significant difference between the untreated and treated groups. For IL-8 (Fig. 4O), disease-like monoculture exhibited higher levels after treatment, increasing from 575 pg/mL in untreated cultures to 675 pg/mL following tanezumab exposure (*p* < 0.05). As with IL-1β, healthy monoculture showed no detectable change in response to treatment.

Other analytes, including MMP family members, TIMPs, and inflammatory cytokines such as IL-1α, IL-4, IL-6, IL-10, IL-13, MCP-1, TNF-α, and IFN-γ, remained largely unchanged between untreated and tanezumab-treated monocultures under both healthy and disease-like conditions.

The overall similarity between untreated and tanezumab-treated monocultures is further illustrated in Fig. 5, which summarizes pairwise comparisons across all analytes. As shown in Fig. 5A, markers such as MMP-1, MMP-3, IL-6, and IL-8 clustered closely together along both the fold-change and significance axes, indicating broadly similar expression profiles between untreated and treated disease-like monocultures. Although additional comparisons were also generated, including tanezumab versus untreated healthy monoculture and disease versus healthy monoculture under tanezumab exposure, the primary comparison aligned with the study objective was the direct contrast between untreated and treated monoculture groups.

**Figure 5.**
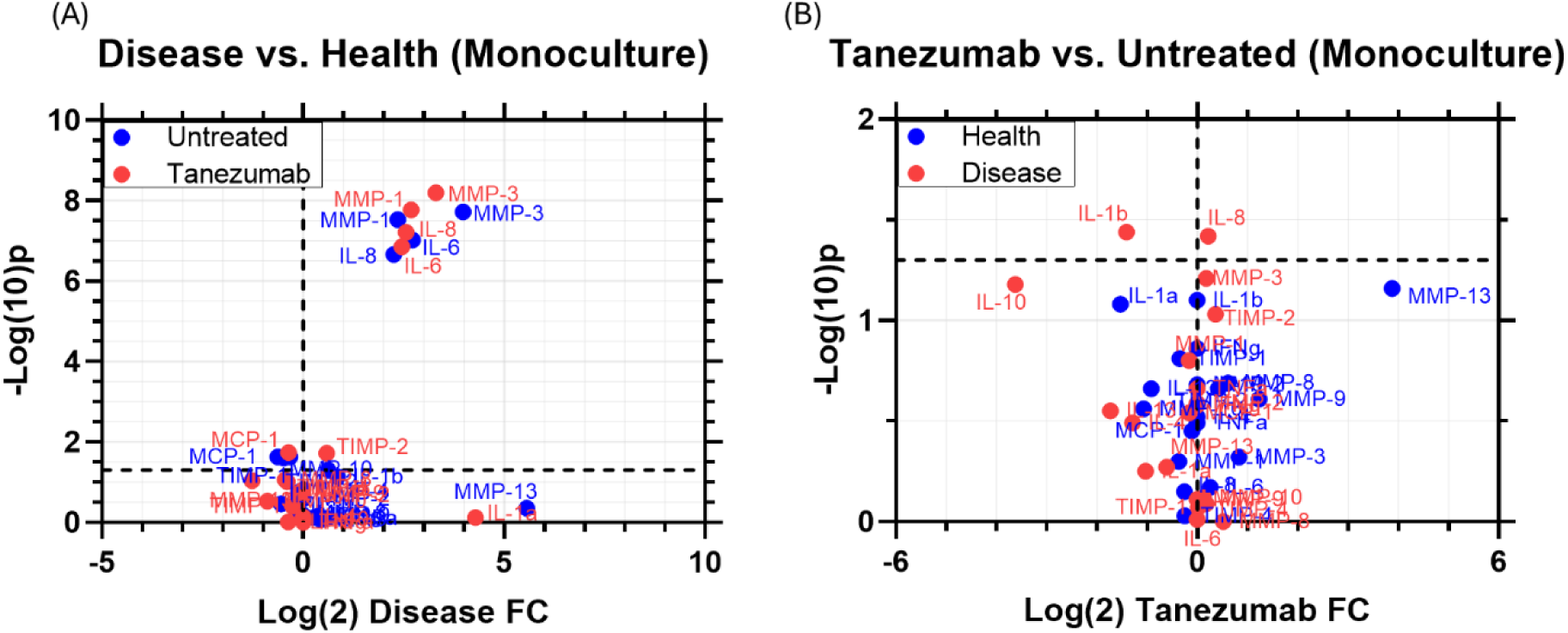
Volcano plots illustrating marker expression changes in human chondrocyte monocultures. **(A)** Disease vs. Health (Monoculture). Volcano plot showing log₂ fold change and -log₁₀(p-value) for inflammatory and matrix-remodeling markers comparing IL-1β-stimulated (disease-like) monocultures to healthy monocultures. Both untreated (blue) and tanezumab-treated (red) groups are displayed, with several markers (e.g., MMP-1, MMP-3, IL-6, IL-8) clustering closely between treatment conditions. **(B)** Tanezumab vs. Untreated (Monoculture). Volcano plot illustrating marker responses to tanezumab within healthy and disease-like monocultures. Marker positions show the fold-change and significance distribution relative to untreated controls under each inflammatory state.

### 4.2 Co-Culture Response to Tanezumab

As shown in Fig. 6, comparison between vehicle (untreated) and tanezumab-treated co-cultures revealed distinct biomarker response patterns across healthy and disease-like conditions For MMP-1 (Fig. 6A), healthy co-culture exhibited higher concentrations after tanezumab exposure, increasing from approximately 4.20 × 10^4^ pg/mL in untreated cultures to approximately 6.20 × 10^4^ pg/mL after treatment (*p* < 0.05). In contrast, disease-like co-culture, which showed higher baseline levels at approximately 7.00 × 10^4^ pg/mL, decreased to approximately 6.00 × 10^4^ pg/mL following treatment. A similar pattern was observed for MMP-3 (Fig. 6C), where healthy co-culture increased from nearly 8.00 × 10^4^ pg/mL to approximately 1.20 × 10^5^ pg/mL after tanezumab exposure (*p* < 0.05), while disease-like co-culture decreased from approximately 1.40 × 10^5^ pg/mL to 1.23 × 10^5^ pg/mL following treatment.

**Figure 6.**
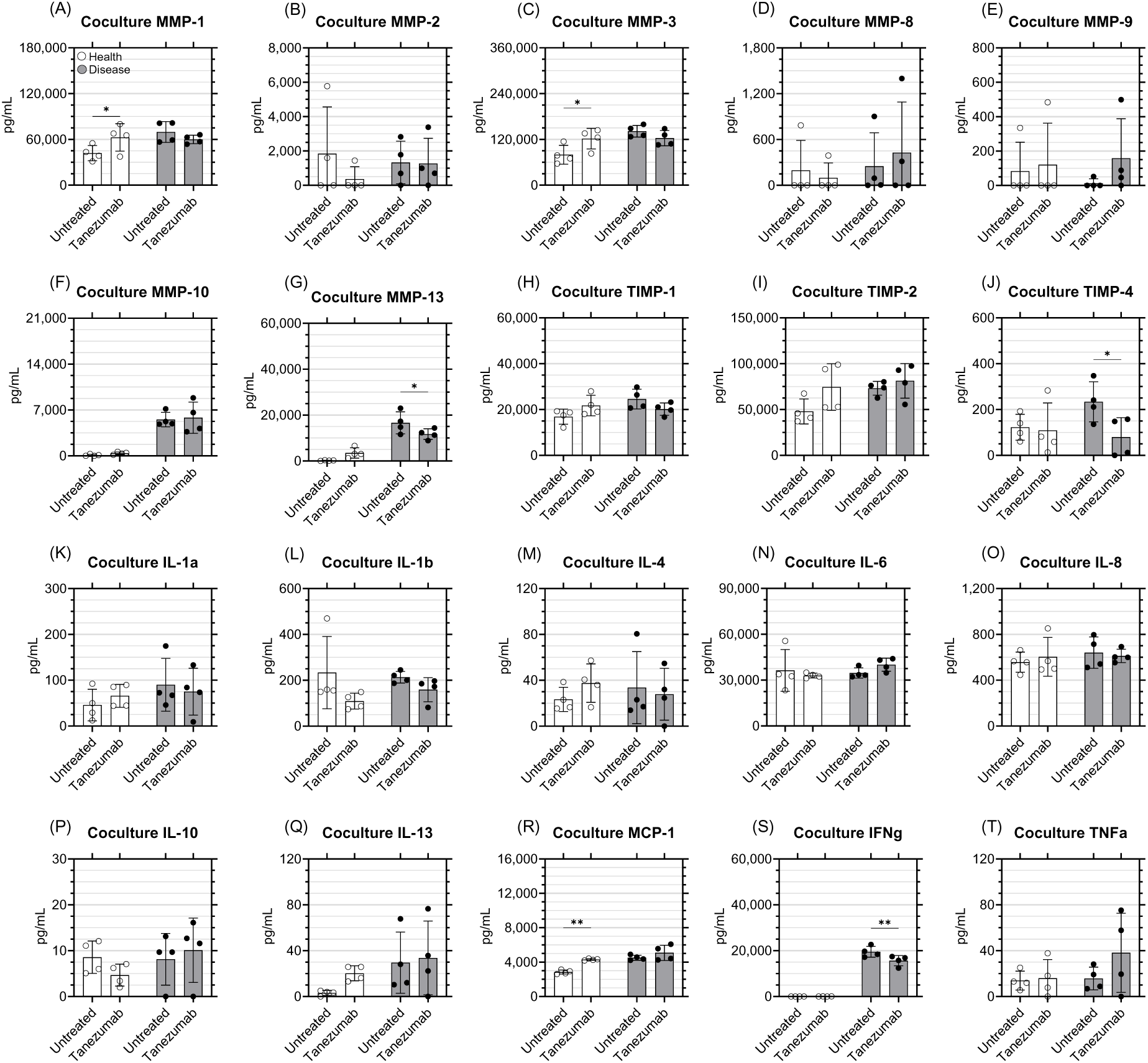
Quantitative secretion profiles of inflammatory and matrix-remodeling markers in microfluidic co-culture models before and after tanezumab exposure. Multiplex ELISA measurements of chondrocyte-derived conditioned media (mean ± SD, n = 4). Statistical significance between untreated and tanezumab-treated conditions is indicated as follows: *p < 0.05; **p < 0.01; ***p < 0.001; ****p < 0.0001. Only a limited subset of analytes demonstrated significant treatment-related changes, whereas most cytokines, chemokines, MMPs, and TIMPs showed no statistically significant differences between conditions

For MMP-13 (Fig. 6G), healthy co-culture exhibited low baseline levels, with untreated samples averaging 172 pg/mL. Following tanezumab exposure, concentrations increased to approximately 3.48 × 10^3^ pg/mL, although this change did not reach statistical significance. In contrast, disease-like co-culture showed substantially higher baseline MMP-13 levels, measuring approximately 1.66 × 10^4^ pg/mL in untreated samples, and decreased to approximately 1.17 × 10^4^ pg/mL after treatment (*p* < 0.05).

For TIMP-4 (Fig. 6J), healthy co-culture remained largely comparable before and after treatment, measuring 123 pg/mL in untreated cultures and 110 pg/mL after tanezumab exposure. In disease-like co-culture, baseline TIMP-4 concentration was 234 pg/mL and decreased to 79.6 pg/mL following treatment (*p* < 0.05).

A similar condition-dependent pattern was observed for MCP-1 and IFN-γ. For MCP-1 (Fig. 6R), healthy co-culture increased from 2.85 × 10^3^ pg/mL in untreated samples to 4.30 × 10^3^ pg/mL after tanezumab treatment (*p* < 0.01). In disease-like co-culture, MCP-1 levels were higher at baseline, measuring 4.52 × 10^3^ pg/mL, and increased slightly to 5.08 × 10^3^ pg/mL after treatment, though this difference was not statistically significant. For IFN-γ (Fig. 6S), healthy co-culture remained low and showed limited variation, measuring 11.8 pg/mL in untreated cultures and 37.4 pg/mL after tanezumab exposure. In contrast, disease-like co-culture showed a markedly higher baseline IFN-γ concentration of 1.95 × 10^4^ pg/mL, which decreased to approximately 1.56 × 10^4^ pg/mL following treatment (*p* < 0.01).

Other analytes, including IL-1α, IL-4, IL-6, IL-8, IL-10, IL-13, TNF-α, MMP-2, MMP-8, MMP-9, MMP-10, TIMP-1, and the remaining measured markers, remained largely unchanged between untreated and tanezumab-treated co-cultures under both healthy and disease-like conditions.

The overall influence of tanezumab across co-culture conditions is further illustrated in Fig. 7, which summarizes the distributions of fold change and significance for healthy and disease-like co-cultures. These volcano plots consolidate global response patterns across all measured analytes and highlight the subset of markers that exhibit measurable shifts under tanezumab treatment. In addition, the number and diversity of OA-relevant analytes represented across the plots may also be considered an important visual feature for interpreting overall response patterns.

**Figure 7.**
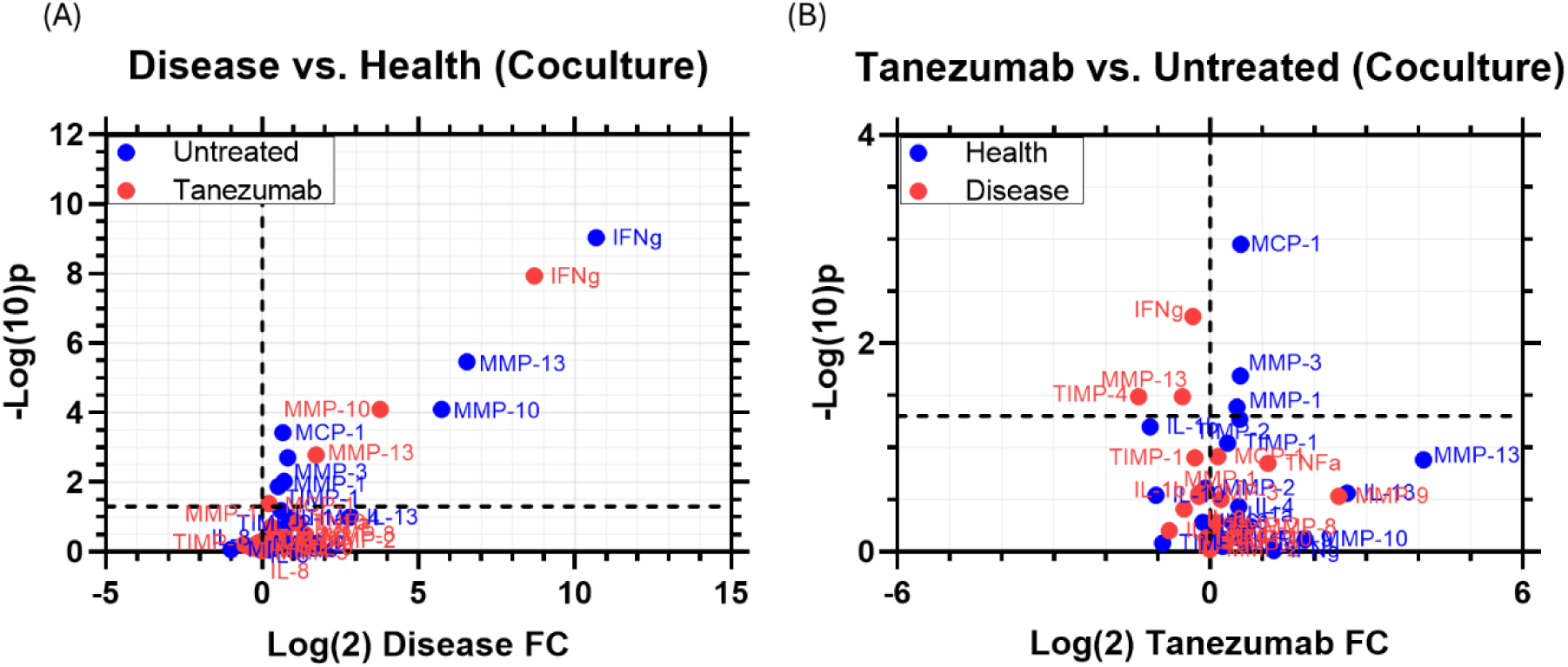
Volcano plot summary of biomarker responses to tanezumab in microfluidic co-culture models. (A) Volcano plot comparing tanezumab-treated vs. untreated co-culture under healthy inflammatory conditions. (B) Volcano plot comparing disease-level vs. healthy co-culture responses in the presence of tanezumab. Each point represents an individual cytokine, chemokine, or metalloproteinase. The x-axis indicates log₂ fold-change, and the y-axis shows -log₁₀(p-value). Horizontal dashed lines denote significance thresholds, and vertical dashed lines denote fold-change reference. These plots provide an integrated visualization of the magnitude and statistical significance of biomarker shifts across co-culture conditions following tanezumab exposure.

## 5 Discussion

This study builds directly on the companion platform-validation manuscript mentioned above, which evaluates the baseline inflammatory and matrix-remodeling behavior of the same M0- and M1-based co-culture configurations. (Mirazi & Wood, 2026) Here, those configurations were used as a treatment-response framework to assess whether multicellular joint crosstalk improves the detection of biomarker changes associated with tanezumab exposure. In this context, the study provides a retrospective evaluation of the platform’s potential predictivity within a defined microphysiological system context of use: identifying early tissue-level inflammatory and safety-relevant biomarker responses to joint-injectable drug exposure that may be missed in chondrocyte monoculture.

Chondrocyte monoculture exhibited minimal and inconsistent molecular responses to tanezumab, indicating that isolated cartilage cells had limited sensitivity for detecting early drug-associated risk signals (Fig. 4 and Fig. 5B). Only IL-1β and IL-8 differed significantly in the disease-like monoculture condition, while matrix-remodeling markers, including MMP-1, MMP-3, and MMP-13, remained largely unchanged. This limited response is consistent with the broader limitation of monoculture systems, which do not capture immune-stromal regulation of cartilage catabolism, inflammatory signaling, or paracrine feedback among joint-associated cell populations. (Deszcz & Bar, 2025; Mehta et al., 2019; Muenzebrock et al., 2022; Pretzel et al., 2009; Tassey et al., 2021) In particular, the absence of macrophage- and stromal-derived cues removes key regulatory components of the joint microenvironment that may be required to reveal more complex responses to pathway-targeted biologics.

In contrast, the healthy (i.e., low-inflammation) M0-based co-culture exhibited clearer, more coordinated molecular responses to tanezumab. Within this low-inflammation environment, in which M0 macrophages, chondrocytes, fibroblast-like cells, and osteoblasts maintained paracrine communication, tanezumab significantly increased MMP-1, MMP-3, and MCP-1 levels relative to untreated controls (Fig. 6A, C, R). An increase in MMP-13 was also observed, although this change did not reach statistical significance (Fig. 6G and Fig. 7B). These coordinated shifts suggest that intact multicellular communication can reveal early alterations in matrix-remodeling and inflammatory signaling that were not apparent in chondrocyte monoculture. This finding is consistent with the established view of OA as a chronic whole-joint disease with a low-grade inflammatory component, in which macrophage-mediated paracrine signaling contributes to chondrocyte catabolic activity, stromal activation, and matrix remodeling. (Berenbaum, 2013; Chen et al., 2020; Cifuentes et al., 2025; De Roover et al., 2023; Loeser et al., 2012; Thomson & Hilkens, 2021; Wu et al., 2020; Zhang et al., 2020)

The M1-based disease-like co-culture produced a markedly different response profile. Because this configuration relied on LPS and IFN-γ stimulation to induce and maintain the M1 phenotype, the system operated in a pro-inflammatory state characterized by elevated baseline levels of cytokines and matrix-remodeling enzymes, as described in detail in the companion manuscript. (Mirazi & Wood, 2026) Under these conditions, most OA-related markers showed limited responsiveness to tanezumab. Notably, decreases in selected biomarkers, including MMP-13 and IFN-γ, were observed following treatment (Fig. 6 and Fig. 7B). Interpreted in isolation, such decreases could be mistaken for beneficial drug effects, despite tanezumab’s unfavorable clinical safety profile. These outcomes may reflect signal compression, a phenomenon observed in hyperinflammatory in vitro systems in which excessive baseline activation restricts the dynamic range available to detect additional drug-induced responses. (Roser et al., 2023) Thus, while the M1 configuration may reproduce aspects of advanced inflammatory activation, it may be less suitable for detecting subtle early safety-relevant perturbations than a lower-inflammation co-culture state.

The differential responses observed among monoculture, M0-based co-culture, and M1-based co-culture also support the broader concept that tanezumab-related effects may depend on the cellular and inflammatory context of the joint. The present model was designed to evaluate tanezumab in an unsupplemented multicellular environment rather than adding exogenous NGF. Although NGF concentrations were not directly measured in this study, prior work supports the plausibility that the included cell populations could generate an endogenous NGF background. NGF is expressed or produced by chondrocytes, synovial fibroblasts, and macrophages, particularly under inflammatory conditions, and osteoblast-lineage cells have also been shown to produce NGF in bone-related contexts. (Pecchi et al., 2014; Shang et al., 2017; Takano et al., 2017) NGF is also upregulated in osteoarthritic joints and regulated by inflammatory signals, with increased NGF expression observed in human subchondral osteochondral tissues, particularly in regions with bone marrow lesions. (Aso et al., 2024; Pecchi et al., 2014; Zhao et al., 2024) In human knee OA specimens, increased NGF expression has been observed in subchondral osteochondral tissues, particularly in regions with bone marrow lesions. (Aso et al., 2024) In osteoarthritic mice, synovial fibroblasts and macrophages have also been shown to produce NGF in response to IL-1β and TNF-α. (Takano et al., 2016) Together, these findings support the rationale for testing NGF inhibition in a multicellular system capable of generating local paracrine signals and identify direct quantification of NGF as a relevant future addition.

A major strength of this work is the use of a novel human joint-on-a-chip model that integrates chondrocytes, fibroblasts, osteoblasts, and macrophages within a shared microenvironment, enabling joint-level paracrine signaling that cannot be recapitulated in monoculture. The paired M0 and M1 configurations, operated under matched experimental conditions, allow direct comparison of how the inflammatory state influences the detection of tanezumab-related molecular changes. These features help define a potential context of use for the platform as an early, human-relevant biomarker response system to evaluate whether joint-targeting (e.g., injectable) therapies affect multicellular inflammatory and joint-safety-relevant matrix remodeling networks. The present findings are consistent with the possibility that NGF blockade may alter local joint-cell signaling, rather than only interrupting sensory neuron activation. More broadly, they suggest that low-inflammation multicellular models may be particularly useful for detecting early tissue-specific safety perturbations that could be masked in strongly stimulated disease models or missed entirely in monoculture.

Several limitations remain relative to the proposed context of use. The M1 phenotype was generated with IFN-γ and LPS, which reliably induce classical macrophage activation but may not fully reflect sterile, OA-relevant inflammation. The platform also focuses on soluble paracrine signaling without incorporating mechanical loading, quantification of actual ECM remodeling, or neurovascular influences. Additionally, NGF levels and tanezumab target engagement were not directly measured, limiting interpretation of the specific pathway interactions responsible for the observed biomarker changes. Moreover, the study captures short-term responses, whereas tanezumab’s clinical adverse events developed over longer periods of exposure.

Future work could integrate OA-relevant macrophage cues, extended or repeated dosing, mechanical stimulation, or additional joint-resident cells such as primary synoviocytes, adipocytes, or sensory neurons. Combining transcriptomic profiling with quantitative proteomic analyses may also help clarify the mechanisms driving the distinct responses across inflammatory states. Together, these extensions would strengthen the use of multicellular joint-on-a-chip systems for identifying tissue-level drug responses that may be missed by conventional monoculture assays.

## 6 Conclusions

Overall, these findings suggest that incorporating human chondrocytes, osteoblasts, fibroblasts, and macrophages within an integrated microfluidic co-culture system may improve the detection of early drug-induced responses that are not readily apparent in simpler monoculture models. In this study, human chondrocyte monoculture showed limited biomarker responsiveness to tanezumab, whereas the co-culture system, particularly the healthy model (low-grade inflammation/M0), revealed clearer changes in selected catabolic and inflammatory markers that may be more relevant to the drug’s known clinical safety concerns.

By contrast, the disease model (high-grade inflammation/M1) showed more muted or less distinct response patterns, suggesting that a strongly inflamed baseline may reduce the visibility of early mechanistic signals. Together, these observations indicate that both multicellular communication and baseline inflammatory context may influence how clearly drug-related effects can be detected in osteoarthritis models.

Taken together, this microphysiological joint model may provide a useful, human-relevant platform to support a context of use for improving the early assessment of potential safety-related biomarker responses in the preclinical evaluation of OA therapeutics.

## Acknowledgments and Disclaimers

## Acknowledgments

This material is based upon work supported by the National Science Foundation under Grant No. 2517512 (2234590).

## Ethics Statement

The human osteoblasts, chondrocytes, fibroblasts, and macrophages used in this study were obtained from commercial sources (PromoCell and ATCC). All cell sources were anonymized and supplied with verified informed consent and ethical clearance from the vendors; therefore, no additional institutional ethical approval was required for this work.

## Data Availability

Data supporting the findings of this study are available from the corresponding author upon reasonable request.

## Use of Generative Artificial Intelligence

A generative artificial intelligence (AI) tool, ChatGPT (OpenAI, GPT-5.3), was used in a limited capacity to enhance language quality and readability during the preparation of this manuscript. All scientific content, data interpretation, experimental design, and conclusions were independently conceived, developed, and rigorously verified by the authors. The authors retain full responsibility for the accuracy, integrity, and originality of the work.

## Author Contributions

HM: Conceptualization, Data Curation, Formal Analysis, Investigation, Methodology, Software, Validation, Visualization, Writing - original draft, Writing - review and editing. SW: Conceptualization, Data Curation, Formal Analysis, Funding Acquisition, Investigation, Methodology, Project Administration, Resources, Software, Supervision, Validation, Visualization, Writing - original draft, Writing - review and editing.

## References

Aso, K., Sugimura, N., Wada, H., Deguchi, S., & Ikeuchi, M. (2024). Increased nerve growth factor expression and osteoclast density are associated with subchondral bone marrow lesions in osteoarthritic knees. Osteoarthritis and Cartilage Open, 6(3), 100504. 10.1016/j.ocarto.2024.100504

Berenbaum, F. (2013). Osteoarthritis as an inflammatory disease (osteoarthritis is not osteoarthrosis!). Osteoarthritis and Cartilage, 21(1), 16–21. 10.1016/j.joca.2012.11.012

Chen, Y., Jiang, W., Yong, H., He, M., Yang, Y., Deng, Z., & Li, Y. (2020). Macrophages in osteoarthritis: pathophysiology and therapeutics. Am J Transl Res, 12(1), 261–268.

Cifuentes, M., Verdejo, H. E., Castro, P. F., Corvalan, A. H., Ferreccio, C., Quest, A. F. G., Kogan, M. J., & Lavandero, S. (2025). Low-Grade Chronic Inflammation: a Shared Mechanism for Chronic Diseases. Physiology, *40*(1), 4-25. 10.1152/physiol.00021.2024

Danehy, S. (2017). Pfizer and Lilly Receive FDA Fast Track Designation for Tanezumab. https://www.pfizer.com/news/press-release/press-release-detail/pfizer_and_lilly_receive_fda_fast_track_designation_for_tanezumab

De Roover, A., Escribano-Núñez, A., Monteagudo, S., & Lories, R. (2023). Fundamentals of osteoarthritis: Inflammatory mediators in osteoarthritis. Osteoarthritis and Cartilage, 31(10), 1303–1311. 10.1016/j.joca.2023.06.005

Deszcz, I., & Bar, J. (2025). Co-Culture Approaches in Cartilage and Bone Tissue Regeneration. International Journal of Molecular Sciences, 26(12), 5711. https://www.mdpi.com/1422-0067/26/12/5711

Dietz, B. W., Nakamura, M. C., Bell, M. T., & Lane, N. E. (2021). Targeting Nerve Growth Factor for Pain Management in Osteoarthritis—Clinical Efficacy and Safety. Rheumatic Disease Clinics of North America, 47(2), 181–195. 10.1016/j.rdc.2020.12.003

Global, regional, and national burden of osteoarthritis, 1990-2020 and projections to 2050: a systematic analysis for the Global Burden of Disease Study 2021. (2023). Lancet Rheumatol, *5*(9), e508-e522. 10.1016/s2665-9913(23)00163-7

Hochberg, M. C. (2015). Serious joint-related adverse events in randomized controlled trials of anti-nerve growth factor monoclonal antibodies. Osteoarthritis Cartilage, 23 *Suppl 1*, S18–21. 10.1016/j.joca.2014.10.005

Hochberg, M. C., Tive, L. A., Abramson, S. B., Vignon, E., Verburg, K. M., West, C. R., Smith, M. D., & Hungerford, D. S. (2016). When Is Osteonecrosis Not Osteonecrosis?: Adjudication of Reported Serious Adverse Joint Events in the Tanezumab Clinical Development Program. Arthritis Rheumatol, 68(2), 382–391. 10.1002/art.39492

Iannone, F., De Bari, C., Dell’Accio, F., Covelli, M., Patella, V., Lo Bianco, G., & Lapadula, G. (2002). Increased expression of nerve growth factor (NGF) and high affinity NGF receptor (p140 TrkA) in human osteoarthritic chondrocytes. Rheumatology (Oxford), 41(12), 1413–1418. 10.1093/rheumatology/41.12.1413

LaBranche, T. P., Bendele, A. M., Omura, B. C., Gropp, K. E., Hurst, S. I., Bagi, C. M., Cummings, T. R., Grantham, L. E., 2nd, Shelton, D. L., & Zorbas, M. A. (2017). Nerve growth factor inhibition with tanezumab influences weight-bearing and subsequent cartilage damage in the rat medial meniscal tear model. Ann Rheum Dis, *76*(1), 295-302. 10.1136/annrheumdis-2015-208913

Lane, N. E., Schnitzer, T. J., Birbara, C. A., Mokhtarani, M., Shelton, D. L., Smith, M. D., & Brown, M. T. (2010). Tanezumab for the Treatment of Pain from Osteoarthritis of the Knee. New England Journal of Medicine, 363(16), 1521–1531. 10.1056/NEJMoa0901510

Li, S., Cao, P., Chen, T., & Ding, C. (2023). Latest insights in disease-modifying osteoarthritis drugs development. Ther Adv Musculoskelet Dis, 15, 1759720x231169839. 10.1177/1759720x231169839

Loeser, R. F., Goldring, S. R., Scanzello, C. R., & Goldring, M. B. (2012). Osteoarthritis: a disease of the joint as an organ. Arthritis Rheum, 64(6), 1697–1707. 10.1002/art.34453

May, B. (2021). FDA Cites Safety Concerns Over Pfizer and Lilly’s Osteoarthritis Drug Tanezumab. BioSpace. https://www.biospace.com/fda-cites-safety-concerns-over-pfizer-and-lilly-s-osteoarthritis-drug-tanezumab?keywords=Tanezumab%20

McKenzie, H. (2021). Eli Lilly and Pfizer Put Osteoarthritis Pain Drug Tanezumab Out of its Misery. https://www.biospace.com/eli-lilly-and-pfizer-put-once-promising-osteoarthritis-pain-drug-out-of-its-misery

McMahon, S. B., Bennett, D. L. H., Priestley, J. V., & Shelton, D. L. (1995). The biological effects of endogenous nerve growth factor on adult sensory neurons revealed by a trkA-IgG fusion molecule. Nature Medicine, 1(8), 774–780. 10.1038/nm0895-774

Mehta, S., Akhtar, S., Porter, R. M., Önnerfjord, P., & Bajpayee, A. G. (2019). Interleukin-1 receptor antagonist (IL-1Ra) is more effective in suppressing cytokine-induced catabolism in cartilage-synovium co-culture than in cartilage monoculture. Arthritis Research & Therapy, 21(1), 238. 10.1186/s13075-019-2003-y

Mirazi, H., & Wood, S. T. (2025). Macrophage-Driven Inflammatory Crosstalk in a Human Joint-on-a-Chip Model of Osteoarthritis. Front Pharmacol, 16, 1579228. 10.3389/fphar.2025.1579228

Mirazi, H., & Wood, S. T. (2026). Microfluidic Osteoarthritis-on-a-Chip: Modeling Human Joint Inflammation. Manuscript submitted concurrently for publication in Biofabrication, 2026.2002.2006.704398. 10.64898/2026.02.06.704398

Muenzebrock, K. A., Kersten, V., Alblas, J., Garcia, J. P., & Creemers, L. B. (2022). The Added Value of the “Co” in Co-Culture Systems in Research on Osteoarthritis Pathology and Treatment Development [Review]. Frontiers in Bioengineering and Biotechnology, Volume 10 - 2022. 10.3389/fbioe.2022.843056

Oo, W. M., & Hunter, D. J. (2022). Repurposed and investigational disease-modifying drugs in osteoarthritis (DMOADs). Therapeutic Advances in Musculoskeletal Disease, 14, 1759720X221090297. 10.1177/1759720X221090297

Pecchi, E., Priam, S., Gosset, M., Pigenet, A., Sudre, L., Laiguillon, M.-C., Berenbaum, F., & Houard, X. (2014). Induction of nerve growth factor expression and release by mechanical and inflammatory stimuli in chondrocytes: possible involvement in osteoarthritis pain. Arthritis Research & Therapy, 16(1), R16. 10.1186/ar4443

Pfizer. (2010, June 22, 2010). Pfizer Suspends Tanezumab Osteoarthritis Clinical Trial Program. Retrieved June 17, 2026 from https://www.pfizer.com/news/press-release/press-release-detail/pfizer_suspends_tanezumab_osteoarthritis_clinical_trial_program

A PHASE 3, MULTICENTER, RANDOMIZED, LONG TERM STUDY OF THE SAFETY OF TANEZUMAB IN PATIENTS WITH OSTEOARTHRITIS OF THE KNEE OR HIP. (2008). https://clinicaltrials.gov/study/NCT00809783

Pretzel, D., Pohlers, D., Weinert, S., & Kinne, R. W. (2009). In vitro model for the analysis of synovial fibroblast-mediated degradation of intact cartilage. Arthritis Res Ther, 11(1), R25. 10.1186/ar2618

Qvist, P., Bayjensen, A., Christiansen, C., Dam, E., Pastoureau, P., & Karsdal, M. (2008). The disease modifying osteoarthritis drug (DMOAD): Is it in the horizon?☆. Pharmacological Research, 58(1), 1–7. 10.1016/j.phrs.2008.06.001

Raychaudhuri, S. P., & Raychaudhuri, S. K. (2009). The regulatory role of nerve growth factor and its receptor system in fibroblast-like synovial cells. Scand J Rheumatol, 38(3), 207–215. 10.1080/03009740802448866

Roemer, F. W., Hochberg, M. C., Carrino, J. A., Kompel, A. J., Diaz, L., Hayashi, D., Crema, M. D., & Guermazi, A. (2023). Role of imaging for eligibility and safety of a-NGF clinical trials. Ther Adv Musculoskelet Dis, 15, 1759720x231171768. 10.1177/1759720x231171768

Roser, L. A., Luckhardt, S., Ziegler, N., Thomas, D., Wagner, P. V., Damm, G., Scheffschick, A., Hewitt, P., Parnham, M. J., & Schiffmann, S. (2023). Immuno-inflammatory in vitro hepatotoxicity models to assess side effects of biologicals exemplified by aldesleukin [Original Research]. Frontiers in Immunology, Volume 14 - 2023. 10.3389/fimmu.2023.1275368

Schmelz, M., Mantyh, P., Malfait, A. M., Farrar, J., Yaksh, T., Tive, L., & Viktrup, L. (2019). Nerve growth factor antibody for the treatment of osteoarthritis pain and chronic low-back pain: mechanism of action in the context of efficacy and safety. Pain, 160(10), 2210–2220. 10.1097/j.pain.0000000000001625

Shang, X., Wang, Z., & Tao, H. (2017). Mechanism and therapeutic effectiveness of nerve growth factor in osteoarthritis pain. Therapeutics and Clinical Risk Management, 13(null), 951–956. 10.2147/TCRM.S139814

Takano, S., Uchida, K., Inoue, G., Miyagi, M., Aikawa, J., Iwase, D., Iwabuchi, K., Matsumoto, T., Satoh, M., Mukai, M., Minatani, A., & Takaso, M. (2017). Nerve growth factor regulation and production by macrophages in osteoarthritic synovium. Clin Exp Immunol, 190(2), 235–243. 10.1111/cei.13007

Takano, S., Uchida, K., Miyagi, M., Inoue, G., Fujimaki, H., Aikawa, J., Iwase, D., Minatani, A., Iwabuchi, K., & Takaso, M. (2016). Nerve Growth Factor Regulation by TNF-α and IL-1β in Synovial Macrophages and Fibroblasts in Osteoarthritic Mice. J Immunol Res, 2016, 5706359. 10.1155/2016/5706359

Tassey, J., Sarkar, A., Van Handel, B., Lu, J., Lee, S., & Evseenko, D. (2021). A Single-Cell Culture System for Dissecting Microenvironmental Signaling in Development and Disease of Cartilage Tissue. Front Cell Dev Biol, 9, 725854. 10.3389/fcell.2021.725854

Thomson, A., & Hilkens, C. M. U. (2021). Synovial Macrophages in Osteoarthritis: The Key to Understanding Pathogenesis? [Mini Review]. Frontiers in Immunology, Volume 12 - 2021. 10.3389/fimmu.2021.678757

Tiseo, P. J., Kivitz, A. J., Ervin, J. E., Ren, H., & Mellis, S. J. (2014). Fasinumab (REGN475), an antibody against nerve growth factor for the treatment of pain: Results from a double-blind, placebo-controlled exploratory study in osteoarthritis of the knee. Pain, 155(7). https://journals.lww.com/pain/fulltext/2014/07000/fasinumab regn475_,_an_anti body_against_nerve.13.aspx

Wadhwa, N., Crowe, R., Dial, J., & Hern, K. (2019). Pfizer and Lilly Announce Top-Line Results From Second Phase 3 Study of Tanezumab in Osteoarthritis Pain. https://www.pfizer.com/news/press-release/press-release-detail/pfizer_and_lilly_announce_top_line_results_from_second_phase_3_study_of_tanezumab_in_osteoarthritis_pain

Woolf, C. J., Safieh-Garabedian, B., Ma, Q. P., Crilly, P., & Winter, J. (1994). Nerve growth factor contributes to the generation of inflammatory sensory hypersensitivity. Neuroscience, 62(2), 327–331. 10.1016/0306-4522(94)90366-2

Wu, C. L., Harasymowicz, N. S., Klimak, M. A., Collins, K. H., & Guilak, F. (2020). The role of macrophages in osteoarthritis and cartilage repair. Osteoarthritis Cartilage, 28(5), 544–554. 10.1016/j.joca.2019.12.007

Wu, R., Guo, Y., Chen, Y., & Zhang, J. (2025). Osteoarthritis burden and inequality from 1990 to 2021: a systematic analysis for the global burden of disease Study 2021. Scientific Reports, 15(1), 8305. 10.1038/s41598-025-93124-z

Zhang, B., Tian, X., Qu, Z., Liu, J., & Yang, L. (2021). Relative Efficacy and Safety of Tanezumab for Osteoarthritis: A Systematic Review and Meta-analysis of Randomized-Controlled Trials. The Clinical Journal of Pain, 37(12). https://journals.lww.com/clinicalpain/fulltext/2021/12000/relative_efficacy_and_sa fety_of_tanezumab_for.8.aspx

Zhang, H., Cai, D., & Bai, X. (2020). Macrophages regulate the progression of osteoarthritis. Osteoarthritis and Cartilage, 28(5), 555–561. 10.1016/j.joca.2020.01.007

Zhao, D., Luo, M. H., Pan, J. K., Zeng, L. F., Liang, G. H., Han, Y. H., Liu, J., & Yang, W. Y. (2022). Based on minimal clinically important difference values, a moderate dose of tanezumab may be a better option for treating hip or knee osteoarthritis: a meta-analysis of randomized controlled trials. Ther Adv Musculoskelet Dis, 14, 1759720x211067639. 10.1177/1759720x211067639

Zhao, L., Lai, Y., Jiao, H., & Huang, J. (2024). Nerve growth factor receptor limits inflammation to promote remodeling and repair of osteoarthritic joints. Nature Communications, 15(1), 3225. 10.1038/s41467-024-47633-6

